# Bipolar Switching by HCN Voltage Sensor Underlies Hyperpolarization Activation

**DOI:** 10.1101/406231

**Authors:** John Cowgill, Vadim A. Klenchin, Claudia Alvarez-Baron, Debanjan Tewari, Alexander Blair, Baron Chanda

## Abstract

Despite sharing a common architecture with archetypal voltage-gated ion channels (VGIC), the hyperpolarization- and cyclic AMP-activated ion (HCN) channels open upon hyperpolarization rather than depolarization. The basic motions of voltage sensor and pore gates are conserved implying that these domains are inversely coupled in HCN channels. Using structure-guided protein engineering, we systematically assembled an array of mosaic channels that display the full complement of voltage-activation phenotypes observed in the VGIC superfamily. Our studies reveal that the voltage-sensing S3b-S4 transmembrane segment of the HCN channel has an intrinsic ability to drive pore opening in either direction. Specific contacts at the pore-voltage sensor interface and unique interactions near the pore gate forces the HCN channel into a hERG-like inactivated state, thereby obscuring their opening upon depolarization. Our findings reveal an unexpected common principle underpinning voltage gating in the VGIC superfamily and identify the essential determinants of gating polarity.

## INTRODUCTION

A salient feature of a living cell is that its resting membrane potential is more negative inside than outside. Electrical signaling is mediated by voltage-gated ion channels which are activated by membrane depolarization in response to a variety of stimuli (Hodgkin and Huxley, 1952). However, a subclass of these channels which include the closely related hyperpolarization-activated and cyclic nucleotide-gated (HCN) channels are activated by the moving the potential in the negative direction, away from the threshold for generating electrical impulses. These channels play a unique role in setting the “resting” membrane potential in a variety of cells (Ludwig et al., 1998; Santoro et al., 1998). In autonomously rhythmic cells, such as the sinoatrial node, they drive the membrane potential towards threshold and are, therefore, also known as pacemaking channels (Brown et al., 1979). New cryo-EM structures show that despite some clear differences, the overall structures of the human HCN channel and depolarization-activated channels are remarkably similar (Lee and MacKinnon, 2017). This naturally raises questions regarding the nature of structural and molecular determinants of gating polarity in the voltage-gated ion channel superfamily (VGIC).

The HCN channel is one of the most well characterized hyperpolarization-activated ion channels. They belong to cyclic nucleotide-binding domain (CNBD) family of ion channels which also includes depolarization activated EAG (ether-à-go go) and hERG (human ether-à-go go related) channels, and a prototypical cyclic nucleotide-activated (CNG) ion channel (Craven and Zagotta, 2006; James and Zagotta, 2018; Yu et al., 2005). EAG channels behave like canonical depolarization-activated ion channels (Warmke et al., 1991; Wu et al., 1983; Zhong and Wu, 1991) but hERG channels are functionally unique. These channels open on depolarization but their depolarization-dependent opening is obscured by rapid inactivation resulting in very little currents at those potentials. Upon repolarization, they are able to pass large currents because the channel recovers from inactivation rapidly but is slow to close (Sanguinetti et al., 1995; Smith et al., 1996; Trudeau et al., 1995). Thus, from a functional standpoint in a cardiac muscle cell, hERG in a way resembles a hyperpolarization-activated ion channel.

HCN channels which open only upon hyperpolarization are considered mechanistically distinct from hERG for many reasons. First, the activation of these channels is characterized by sigmoidicity suggesting that the opening of these channels involve multiple closed states rather than simply recovering from an inactivated state (Altomare et al., 2001). Second, there is no evidence that further hyperpolarization will close these channels as expected if the hyperpolarized opening was simply recovery from inactivation (Miller and Aldrich, 1996). Third, residue accessibility measurements on the voltage sensor show that the channel opening is associated with downward movement of the S4 (Mannikko et al., 2002), in contrast to hERG inactivation which is associated with pore collapse (Smith et al., 1996). Finally, studies measuring accessibility of pore residues (Rothberg et al., 2002; Rothberg et al., 2003) and blocker-binding sites (Shin et al., 2001) have established that the intracellular pore gates regulate the access to the ion conduction pathway analogous to canonical outwardly rectifying channels. Thus, the emerging consensus is that the voltage sensor movement must be inversely coupled to pore opening in hyperpolarization-activated channels compared to the depolarization-activated counterparts.

The newly available structures of the EAG (Whicher and MacKinnon, 2016), hERG (Wang and MacKinnon, 2017), and HCN (Lee and MacKinnon, 2017) channels provide insights into possible determinants of outward vs. inward rectification. The traditional chimeragenesis approach of identifying the molecular elements responsible for a functional phenotype has been remarkably successful in identifying self-contained catalytic or functional sites but much less useful in reconstructing long range allosteric interaction pathways involved in signal transduction (Heginbotham et al., 1992; Herzberg et al., 1998; Kobilka et al., 1988; Lu et al., 2001; Wainger et al., 2001). Here, we tackled this challenge in two stages. First, we engineered an array of chimeras using closely related inward and outward rectifiers after identifying the junction points for making swaps. Next, we introduced interactions that are likely to be involved in coupling the voltage sensor and pore gates to create “mosaic” channels. Our hierarchical protein engineering approach reveals that the voltage-sensing S3b-S4 segment of HCN channels is a bipolar switch that can drive the pore opening in both hyperpolarized and depolarized direction. In the HCN, the opening at depolarized potentials is abrogated by hERG-like inactivation controlled by other structural elements including key residues near the gate, the S4-S5 interface, and the C-terminal cytoplasmic region. Taken together, our studies reveal a common mechanistic thread that links the voltage-dependent gating properties of all channels in this superfamily.

## RESULTS

### Voltage-Sensing Segment of HCN Can Drive Opening during Both Hyperpolarization and Depolarization

Despite major differences in function, the overall architecture of HCN1 and EAG1 are remarkably similar (Figure 1A) (Lee and MacKinnon, 2017; Whicher and MacKinnon, 2016). Detailed examination reveals that several structural elements including the charge transfer center, pore module, and the cytoplasmic C-terminal region are highly conserved (Figure 1B). Nevertheless, there are key differences between HCN channels and depolarization-activated members of KCNH family. HCN channels contain an N-terminal HCN-specific domain in place of the PAS domain found in the KCNH channels (Cabral et al., 1998; Haitin et al., 2013) and their S4 transmembrane helix is at least two helical turns longer than its counterpart in the EAG1 and hERG (Lee and MacKinnon, 2017).

**Figure 1.**
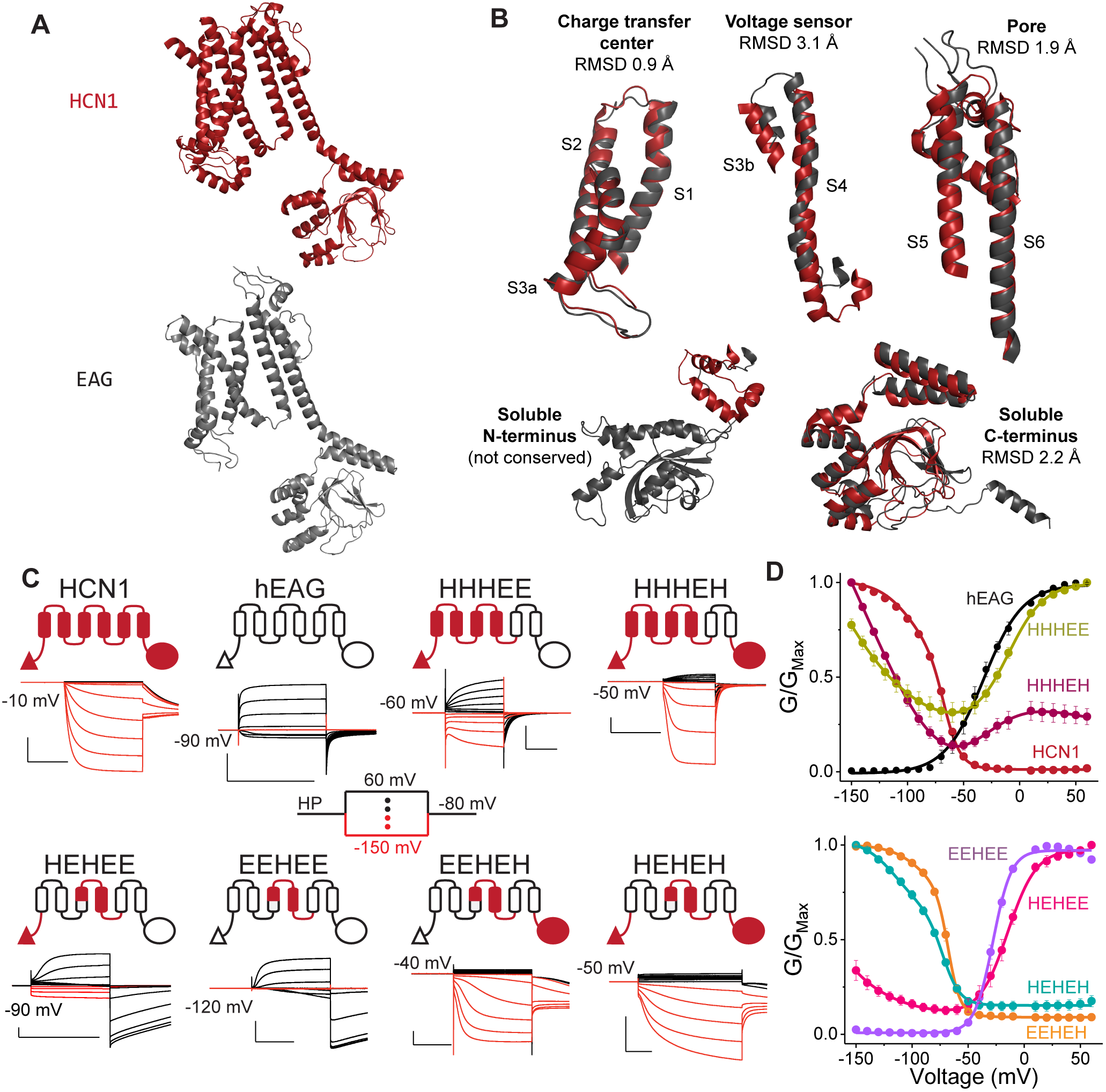
HCN Voltage-sensing Domain is a Bipolar Switch. (A) Monomeric structures of hHCN1 and rEAG1. The color-coding for HCN1 (red) and EAG1 (black) is used throughout all figure panels. The N-terminus of rEAG1 is omitted for clarity. (B) Structural alignment between hHCN1 and rEAG1 for the five modular segments used in generating the chimeric channels. (C) Cartoon representations of chimeric channels and representative traces for currents elicited in cut-open voltage clamp (COVG) under symmetrical solutions (100 mM K^+^_internal_ /100 mM K^+^_external_). Currents in response to depolarizing pulses are colored black and responses to hyperpolarizing pulses are colored red for clarity. Scale bars denote 2 μA (vertical) and 500 ms (horizontal). (D) Conductance-voltage (GV) curves for the parent and chimeric channels. Error bars are SEM from n= 5 (HHHEH and EEHEE) or 6 (HEHEH) or 4 (all others) independent measurements.

To identify the role of each of these structural elements, we used the chimeragenesis approach of swapping these elements between the HCN homologs (mHCN1, mHCN2, and spHCN) and EAG1 channels. The availability of the high-resolution structures for both rEAG1 and hHCN1 (Lee and MacKinnon, 2017; Whicher and MacKinnon, 2016) enables precise structure-aided sequence alignment between the members of this family (Figure S1) and allows identification of the putative junction points that are likely to minimize the perturbation of the local structure. To identify the most optimal junction points, we created multiple chimeras with different junction points (Figure S2) and tested them using electrophysiology. Based on this preliminary scan, we empirically identified four primary junction points that demarcate five structural elements common to these channels (Figure 1C). Henceforth, we will denote these chimeras using a five-letter code representing the origin of each structural element-H for mHCN1 and E for hEAG.

Given the modularity of the voltage-sensing and pore domains (Arrigoni et al., 2013), we first tested the HHHEE chimera that contains the voltage-sensing domain of HCN1 and pore domain of EAG1 (Figure 1D and 1E). Remarkably, this chimera opens in a voltage-dependent manner both at hyperpolarizing and depolarizing potentials while closing at intermediate potentials (HHHEE in Figure 1D). This behavior, which we refer to as Janus phenotype, establishes that the hyperpolarization and depolarization activation pathways are distinct and that the HCN voltage-sensing domain is intrinsically capable of gating the pore in both directions. Swapping in the soluble C-terminus region of the HCN channel into this chimera (HHHEH) maintains the basic phenotype but reduces the depolarization-activated currents. Together, these results suggest that the C-terminus of HCN contributes to shutting the depolarization-activated pathway.

The voltage-sensing domain has three structural elements-the N-terminal HCN domain (Lee and MacKinnon, 2017), the charge transfer center (Tao et al., 2010), and the voltage sensor paddle (Jiang et al., 2003). Replacement of the charge transfer center in HHHEE chimera with its equivalent from EAG1 (HEHEE) results in a channel which primarily activates upon depolarization although these channels can still open upon hyperpolarization. Further replacement of the N-terminus with that of EAG1 (EEHEE chimera) completely abolishes activation by hyperpolarization and this construct behaves like a canonical outward rectifying channel (Figures 1D and 1E). These findings unequivocally establish that the HCN S4 paddle is not sufficient to drive the channels to open upon hyperpolarization and it requires cumulative contribution from both the charge transfer center and N-terminus to stabilize the hyperpolarization-activated opening as predicted based on the structural proximity of these elements (Lee and MacKinnon, 2017).

The role of the HCN C-terminus was further examined by generating two other chimeras EEHEH and HEHEH. Surprisingly, these chimeras show activation upon hyperpolarization but very little activation on depolarization (compare with the EAG-like EEHEE and HEHEE, Figure 1E). Thus, unlike the N-terminus which enhances hyperpolarization-activation without perturbing the depolarization-activation pathway, the C-terminus appears to disfavor activation upon depolarization in addition to enhancing opening at hyperpolarized potentials. Nevertheless, both EEHEH and HEHEH also exhibit substantial voltage-independent leak conductance (~10% of the maximum) indicating that these channels are still missing some elements required for full closure at depolarized potentials.

### Hyperpolarization and Depolarization-activated States Share Same Permeation Pathway

Given the potential for large-scale perturbations in chimeric proteins, it is important to rule out the possibility that the observed currents are simply upregulated endogenous conductances rather than ion flux through the expressed channels. Moreover, subtle perturbations within the voltage-sensing domain have been shown to create additional ion permeation pathways through voltage-sensing domain. These gating pore currents are distinct from the central pore currents (Starace and Bezanilla, 2004; Starace et al., 1997; Wainger et al., 2001). In order to probe the permeation pathway directly, we introduced a cysteine residue near the selectivity filter (hEAG1 mutant A470C, Figure 2A) and evaluated the effect MTSEA modification on ionic currents. The A470C mutants of EAG1 and HHHEH show ~60% reduction in current amplitude upon addition of MTSEA whereas the parent constructs are unaffected (Figures 2B and 2C). Importantly, currents at both hyperpolarizing and depolarizing potentials are inhibited by cysteine modification. These findings establish that the observed conductances, at both potentials, are due to ion flux through the central pore of the expressed channels.

**Figure 2.**
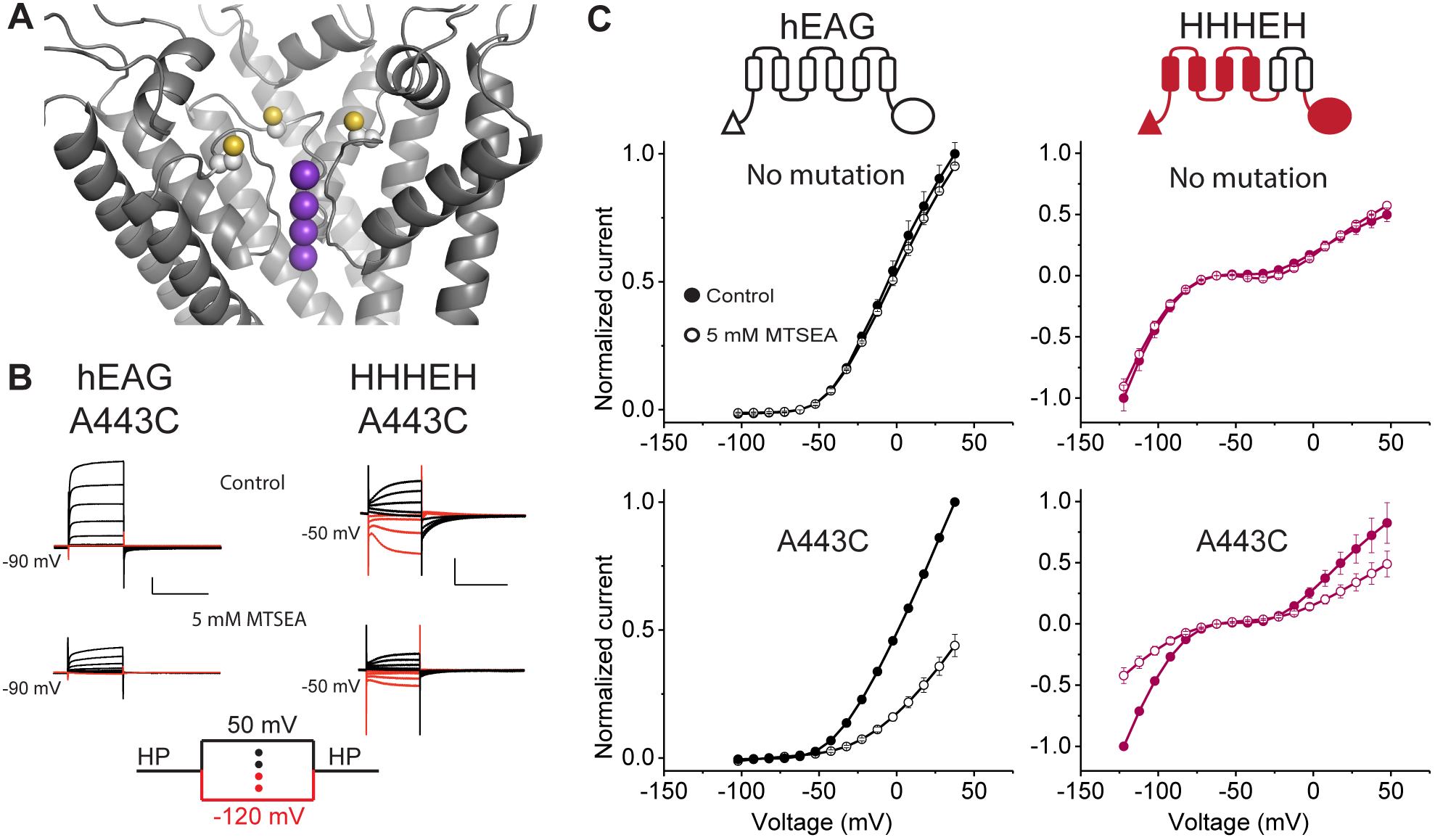
Hyperpolarization- and Depolarization-activated States Share Same Permeation Pathway. (A) Location of the A470C mutation near selectivity filter in the rEAG1 structure (yellow balls for sulfur atoms). Only three subunits are shown for clarity. Purple balls indicate positions of potassium ions as would be expected for EAG1 based on experimentally determined sites in a similar selectivity filter of Kv1.2/2.1. (B) Representative current traces for A470C mutants of EAG1 and HHHEH from two-electrode voltage clamp. Currents from EAG1-A470C were recorded in 100 mM Na^+^/5 mM K^+^ while the HHHEH-A470C was recorded under pseudo-symmetric conditions using the same external solution as in Figure 1. Upper traces represent initial recordings while lower traces represent recordings following application of 5 mM MTSEA. Scale bars denote 2 μA (vertical) and 500 ms (horizontal). (C) Current-voltage relationships for wild type (top) and mutant (bottom) proteins before (filled circles) and after (open circles) MTSEA addition with error bars showing SEM for n= 2 (HHHEH), 3 (EAG1), 6 (EAG1-A470C), or 8 (HHHEH-A470C). See also Figure S3.

To probe the pore gating apparatus, we tested astemizole block. Astemizole is a specific open pore blocker of EAG1 (Garcia-Ferreiro et al., 2004) but does not block currents through HCN1 (Figure S3). We find that this drug blocks outward currents through EEHEE chimera in a manner characteristic of an open pore blocker. Surprisingly, the inward currents through EEHEH or HEHEH chimera are not at all affected by astemizole despite the fact that both of these channels have the EAG1 pore. Astemizole block in both EAG and hERG is highly state-dependent, with only the open and inactive states showing high efficacy binding (Garcia-Ferreiro et al., 2004; Gomez-Varela et al., 2006; Perrin et al., 2008; Zhou et al., 1999). Thus, the inability of astemizole to block hyperpolarization-activated currents in our constructs may suggest a different conformation of the hyperpolarization-activated pore gate compared to that of the depolarization-activated gate.

### HCN1-like S4-S5 Interface Introduces Rapid Inactivation Pathway on Depolarization

As noted by Lee and MacKinnon, one of the most striking features of the HCN1 structure is the tight juxtaposition of the S4 and S5 helices in contrast to the depolarization activated channels like EAG and Kv1.2/2.1 (Lee and MacKinnon, 2017; Long et al., 2005; Whicher and MacKinnon, 2016). A reasonable hypothesis is that the unique hairpin-like structure formed by the S4 and S5 helices of HCN1 contributes to the inverted polarity of voltage-dependent response in the HCN family. To test this idea and recapitulate the S4-S5 interface in our chimeras, six S4-facing residues on the S5 helix of EAG1 were mutated to their HCN1 counterparts (Figure 3A). We refer to these constructs as “mosaics” and henceforth they will be denoted with asterisk. The introduction of the mosaic mutations into the HHHEE, HHHEH and EEHEH chimeras resulted in a virtually complete loss of conductance at depolarizing potentials (the HHHE*E, HHHE*H and EEHE*H constructs, Figures 3B and 3C). These chimeras, intended to reestablish the interaction between the HCN voltage sensor and the pore, reconstituted the HCN gating phenotype at positive voltages.

**Figure 3.**
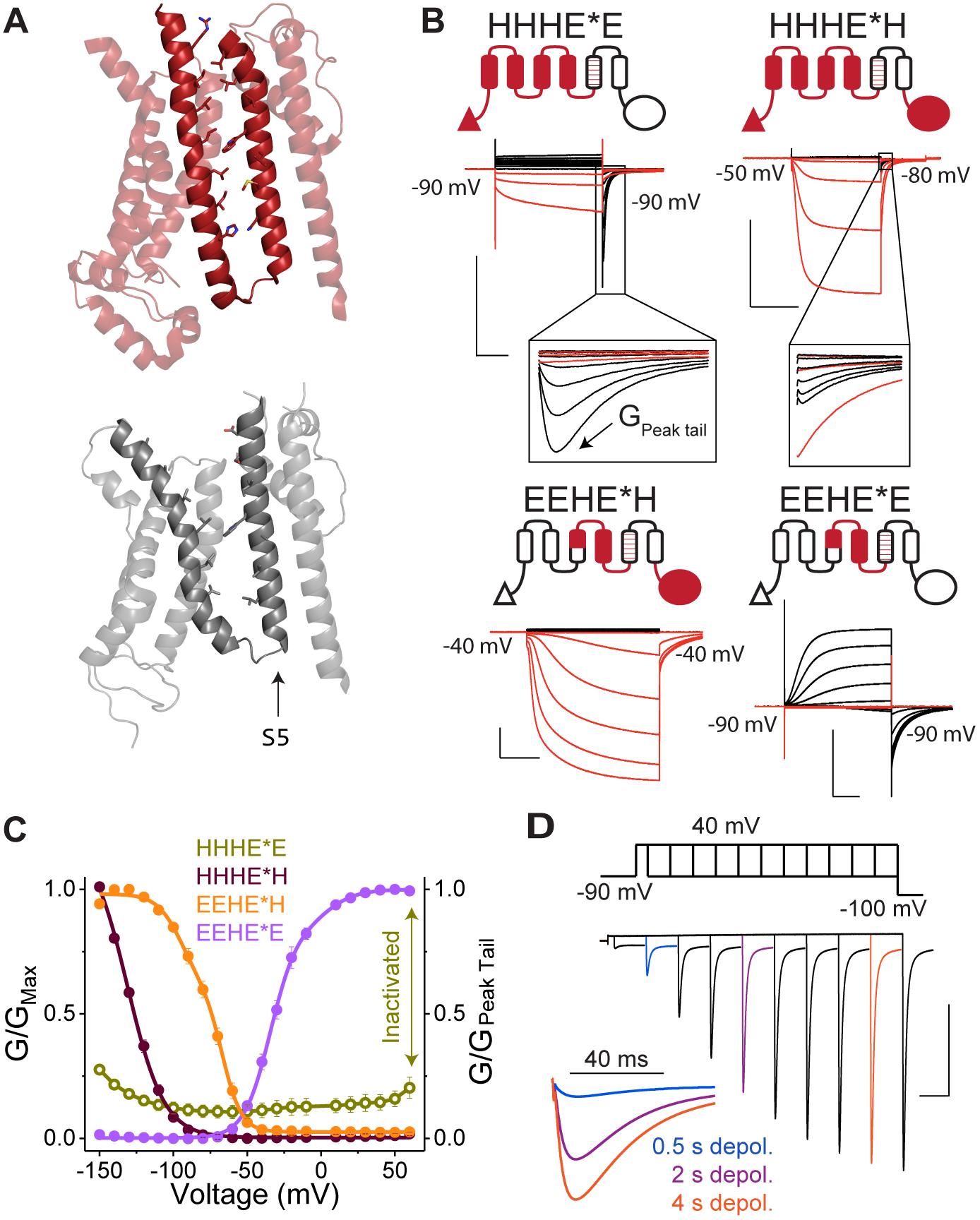
HCN1-like S4-S5 Interface Results in Rapid Inactivation on Depolarization. (A) Structures of the transmembrane regions of hHCN1 (top) and rEAG1 (middle) transmembrane regions highlighting the S4-S5 interface. (B) Cartoon representations of the mosaic chimeras and representative COVG current traces in symmetrical solutions (100 mM K^+^_in_ /100 mM K^+^_ex_). Mosaic mutations are designated with an asterisk in the construct name. Insets show expanded view of tail currents displaying hooked phenotype. (C) Steady state GV curves for mosaic channels depicted in panel B with error bars representing SEM for n= 3 (HHHE*E and EEHE*H) or 4 (all others). Due to the strong inactivation observed in the HHHE*E construct, the steady state conductances were normalized to the maximum tail current observed subsequent to application of depolarizing pulses (labeled as G_peak tail_ in panel B). The data points for HHHE*E are thus plotted as open symbols using a separate axis (right) to emphasize this difference. This convention is used to highlight the fact that only a small fraction of channels are open at steady state. (D) Envelope of tails experimental protocol used with the HHHE*E construct to evaluate tail currents elicited by depolarizing +40 mV pulses of varying duration. Three traces for 0.5, 2, and 4 ms pulses are colored and reproduced at higher temporal resolution on the left. Scale bars in (B) and (D) denote 2 μA (vertical) and 500 ms (horizontal). See also Figure S4.

Interestingly, we observed prominent tail currents upon repolarization in both the HHHE*E and HHHE*H mosaics in addition to enhanced rectification (Figure 3B). These tail currents exhibit a well-defined “hooked” phenotype (Figures 3B), with a peak when the voltage returns to a negative potential. This behavior is characteristic of tail currents in the hERG potassium channel, which recover from an inactivated state before closing following membrane repolarization (Sanguinetti et al., 1995; Trudeau et al., 1995). To test whether the lack of currents on depolarization is due to rapid entry into inactivated state, we used an envelope-of-tails protocol for the HHHE*E (Figure 3C and Figure S4). Under this protocol, the amplitude of peak tail conductance increases over the timescale of seconds with no change to the steady-state conductance (Figure S4). This indicates that the channels undergo rapid inactivation following a slow activation process akin to the hERG channels (Smith et al., 1996; Trudeau et al., 1995). Upon return from depolarized potentials, the channels rapidly recover from inactivation, transiently populating the open state before closing (inset of Figure 3D). These findings establish that the S4-S5 interface of HCN1 plays a critical role in preventing conduction at depolarized potentials through the hERG-like inactivation.

Although addition of the S4-S5 interfacial residues of HCN1 reduced the conductance at depolarized potentials for most mosaics tested, this was not the case for the EEHE*E construct (Figure 3B and 3C). This behavior is not surprising because deletions of soluble N-terminal domain (PAS domain) of EAG1 (Carlson et al., 2013; Terlau et al., 1997) and chimera experiments (Lin et al., 2014) suggest interactions between the PAS domain and cyclic nucleotide-binding homology domain (CNBhD) are necessary to prevent voltage-dependent inactivation in EAG channels. Nevertheless, we cannot rule out the possibility that it is the presence of the HCN1 N-terminus, rather than the absence of PAS domain in the HHHE*E chimera that is required for the rapid inactivation upon depolarization. Unfortunately, this idea cannot be tested directly because deletion of the PAS domain from EEHE*E produces no detectable currents (data not shown).

### Residues at the Tail of S6 Helix Destabilize the Depolarization-activated Open State

A notable difference between HCN1 and HHHE*H constructs is the lack of hooked tails in HCN channels despite overall similarity in voltage-dependent opening upon depolarization (compare Figure 1D and Figure 3B). Hooked tails are observed because the relative rate of recovery from inactivation is comparable to the rate of entry into the closed state (Sanguinetti et al., 1995; Smith et al., 1996; Trudeau et al., 1995). We posit that there are additional pore residues in HCN channels that further destabilize this depolarization-activated open state and accelerate entry into the closed state. We tested this hypothesis with a series of HHHE*H-based constructs in which an increasingly larger proportion of the pore domain is replaced by HCN1-specific residues (Figure 4A, denoted by H2, H3, H4, and H5). As shown in Figures 4B, in all four constructs the substitutions resulted in a loss of hooked tail currents, demonstrating that only eight residues at the tail of the S6 helix (following the “gate” residue Q398 and Q476 of hHCH1 and rEAG1, respectively) are sufficient to suppress the depolarization-dependent opening and make the mosaic constructs functionally equivalent to HCN1 (see Figure 3B for comparison). In addition, these substitutions caused slight shifts in the voltage-dependence of activation upon hyperpolarization although we are not able to quantitate this due to the lack of saturation of these conductances (Figure 4C).

**Figure 4.**
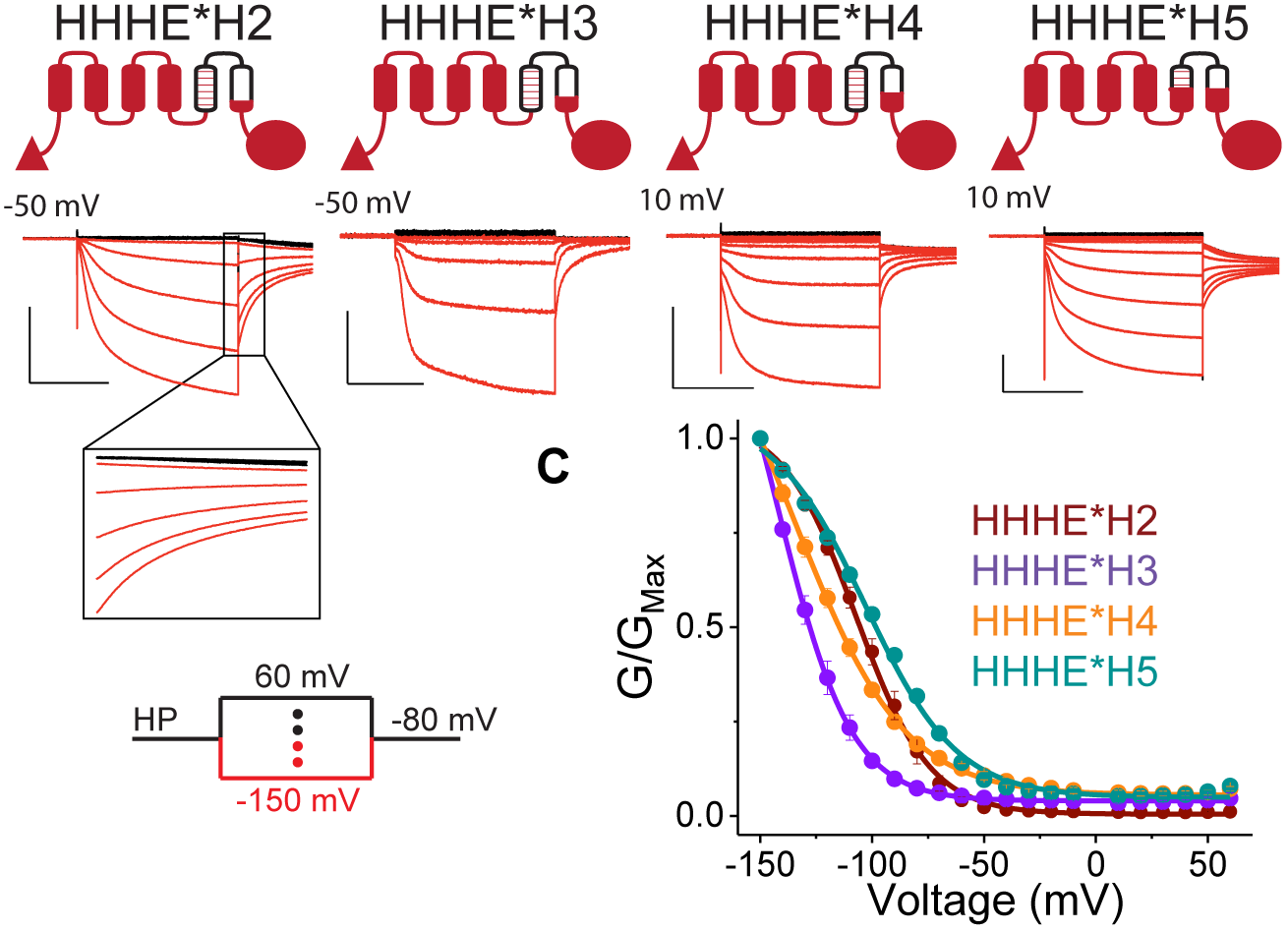
HCN1-specific Residues Proximal to the Gate Contribute to Destabilization of Depolarization-activated Opening. Cartoons and representative COVG currents for the chimeras with various EAG1/HCN1 junction points in S5 and S6. The inset for HHHE*H2 data that clearly indicates a lack of tail currents displaying hooked phenotype. Scale bars denote 2 μA (vertical) and 500 ms (horizontal). (C) GV curves for the new mosaics with error bars showing SEM for n= 3 (HHHE*H4) or 4 (all others). Compare to the parent construct in Figure 3C.

### Role of HCN1 C-terminus in Regulating Polarity of Voltage-dependent Gating

Our data thus far suggest that the C-terminus plays a critical role in regulating the polarity of voltage gating. Replacing the EAG1 C-terminus with its HCN1 counterpart in either the HHHEE or EEHEE chimera (forming the HHHEH and EEHEH chimeras, respectively) leads to a substantial increase in the activation by hyperpolarization compared to depolarization.

Given that voltage-dependent activation of HCN1 is virtually unaffected by removal of its C-terminus (Wainger et al., 2001, see also Figure S5), we speculated that the removal of the EAG1 C-terminus, rather than the addition of the HCN1 equivalent, may account for these observations. However, removal of the C-terminal cytoplasmic region from the HHHE*E mosaic increased currents upon relative to hyperpolarization (Figure 5A and 5B). Thus, the main effect of this truncation is to increase the conductance on depolarization even though the channels are still able to enter the inactivated state rapidly as evidenced by the presence of hooked tail currents.

**Figure 5.**
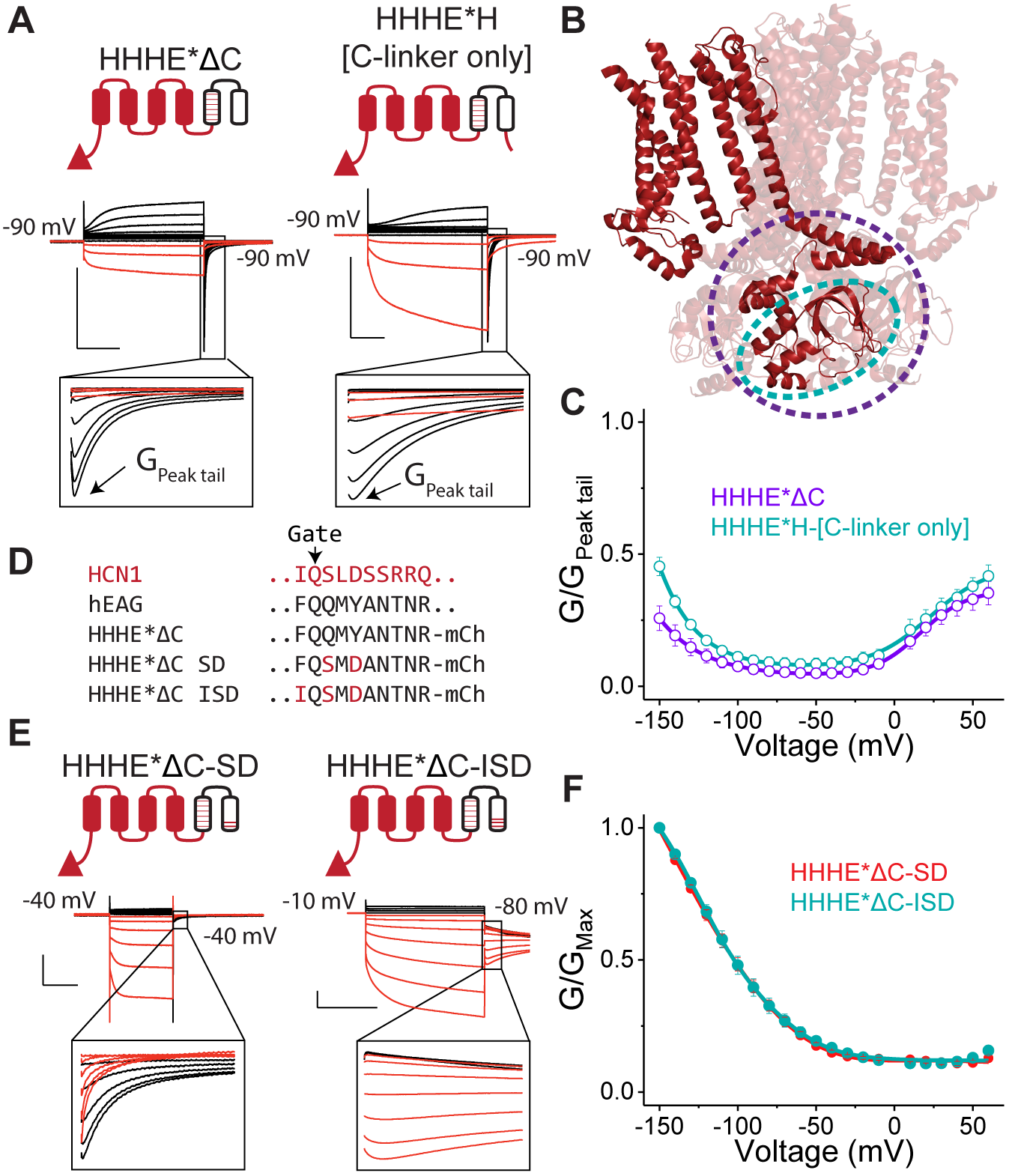
Role of HCN1 C-terminus in Regulating Polarity of Voltage-dependent Gating. (A) Cartoons and representative COVG currents for truncation constructs that lack either the entire cytoplasmic C-terminus (HHHE*ΔC) or only a part of it, retaining R393-S478 of mHCN1 which corresponds to the C-linker (HHHE*ΔC-[C-linker only]). See also Figure S5. (B) Tetrameric structure of hHCN1 with one monomer highlighted. Circles show residues absent in the HHHE*ΔC (purple) and HHHE*HΔC [C-linker only] construct (cyan). (C) GV curves for the C-terminal truncations depicted in (A) with error bars representing SEM for n= 3. Note that, similarly to the treatment of HHHE*E data in Figure 3D, the steady state conductances were normalized to the maximum conductance observed for tail currents. (D) Sequences near the channel’s main gate in S6 helix for mHCN1, hEAG1, HHHE*ΔC and additional mutants made in the HHHE*ΔC background. (E) Cartoons and representative COVG currents for the double and triple mutants of HHHE*ΔC depicted in (D). Note the insets that show that HHHE*ΔC-SD construct still exhibits hooked tail currents on depolarizing pulses while HHHE*ΔC-ISD does not. Scale bars in (A) and (E) denote 2 μA (vertical) 500 ms (horizontal). (F) GV curves for the HHHE*ΔC SD and HHHE*ΔC ISD channels with error bars showing SEM for n= 4 (HHHE*ΔC-SD) or 9 (HHHE*ΔC-ISD). See also Figure S6

As demonstrated by the structure, the longer length of the HCN1 S4-helix enables it to interact with the C-linker (Lee and MacKinnon, 2017). To assess whether this interaction is essential and maintained in the absence of the cyclic nucleotide binding domain (CNBD), we tested mosaic construct that contains the C-linker of HCN1 but lacks the rest of the C-terminus (HHHE*H-[C-linker-only]). The C-linker addition, however, had a minimal impact on voltage-dependent gating (Figures 5A and 5B), indicating that the effect of the HCN1 C-terminus on polarity of gating requires the intact CNBD. Therefore, we conclude that in addition to the C-terminus there are other elements in WT HCN1 that are responsible for preventing reopening upon depolarization.

Owing to the extensive interactions between the C-linker and CNBD, it is still quite likely that the intact CNBD is needed primarily to maintain secondary structure of the C-linker and/or physiologically “proper” tilt of the junction between the C-linker and gate-forming S6 helix. This idea is particularly appealing in the light of the key role that the proximal S6 residues play in preventing activation by depolarization (see HHHE*H2 mosaic in Figure 4). Given that these residues connect directly to the C-linker, we reasoned that they might be equally important even in the absence of the C-terminal fragment. To test this possibility, we replaced the eight EAG1-specific residues at the tip of S6 in the HHHE*ΔC construct with those from HCN1 (note that this construct, HHHE*ΔC2, is a C-terminal deletion of HHHE*H2 mosaic described in the previous section). The resulting channel is primarily activated upon hyperpolarization but also exhibits hooked tail currents upon return to negative membrane potentials from depolarizations (Figure S6). These observations show that the terminal S6 residues are indeed critical and sufficient for disfavoring depolarization activation even in the absence of the C-terminal region. Note, however, that this construct elicits hooked tails upon return from depolarization which is observed neither in the WT HCN nor in HCN1ΔC channels.

Alanine-scanning mutagenesis in this region has shown that two of the S6 tail residues S441 and D443 in mHCN2 (conserved in HCN1 and spHCN) stabilize the closed state (Decher et al., 2004), suggesting that they could be primary determinants of the effects we observe. Substituting these two residues back into the pore in the HHHE*ΔC construct (Q504S and Y506D, hEAG1 numbering) recapitulates the phenotype of the six-residue swap (compare HHHE*ΔC2 in Figure S6A and HHHE*ΔC-SD in Figure 5E). Similarly, the same double mutation introduced into non-mosaic C-terminal truncation construct also displays increased conductance at hyperpolarized compared to depolarized potentials (HHHEΔC and HHHEΔC-SD, Figures S6C and S6D). Examination of the rEAG1 structure reveals that another aromatic residue, F475, immediately precedes the gate and is potentially involved in aromatic stacking with Y479, the equivalent to Y506 in hEAG1 mutated above. The equivalent residue in all HCN-like channels is a highly conserved isoleucine. Substituting this isoleucine into the HHHE*ΔC-SD construct (F502I, hEAG1 numbering) results in no detectable hooked tails upon return from depolarized potentials (compare HHHE*ΔC-SD and HHHE*ΔC-ISD, Figures 5E). Altogether, we can conclude that the three pore mutations are able to functionally substitute for the entire cytoplasmic C-terminus, producing an HCN1-like phenotype as long as the mosaic mutations at the S4-S5 interface are present.

### Long S4 Helix is not Required for Activation by Hyperpolarization

Thus far, all of our chimeras and mosaics that open upon hyperpolarization contain the entire S4 helix of HCN1 which, as revealed by the structure, is unusually long in comparison to the S4 segments of depolarization-activated channels such as hERG and hEAG1 (Lee and MacKinnon, 2017; Wang and MacKinnon, 2017; Whicher and MacKinnon, 2016). Structure-aided sequence alignment shows that spHCN, a sea urchin homolog of HCN1, lacks four residues in this extended region of the S4 (Figures 6A). Using the HEHE*H as a base construct, since it behaves like a prototypical HCN channel, we examined the effect of large deletions inside the S4 segment. Eliminating four residues in the S4-S5 linker, a deletion that must force formation of new S4-S5 helix-turn-helix, and should result in a slight shortening of the S4 helix, results in a channel that retains activation on hyperpolarization with only minor differences compared to the parent construct (Figure 6B). In contrast, removal of eight residues, a deletion that should result in S4 approximately as short as that in EAG1, generates a channel that is still activated on hyperpolarization but undergoes a voltage-dependent inactivation below -100 mV (Figures 6B and 6C). Additionally, this mutant is unable to fully close, with roughly 40% of the peak conductance remaining at depolarizing potentials. These results demonstrate that neither the extra length of the HCN1 S4 nor its residue-specific interactions with the C-linker are the essential determinants of activation by hyperpolarization. The longer S4 may, however, help stabilize the hyperpolarization dependent open state and protect the channel from a voltage-dependent inactivation. Additionally, given the significant basal conductance, it may be important in stabilizing the closed state at depolarized potentials.

**Figure 6.**
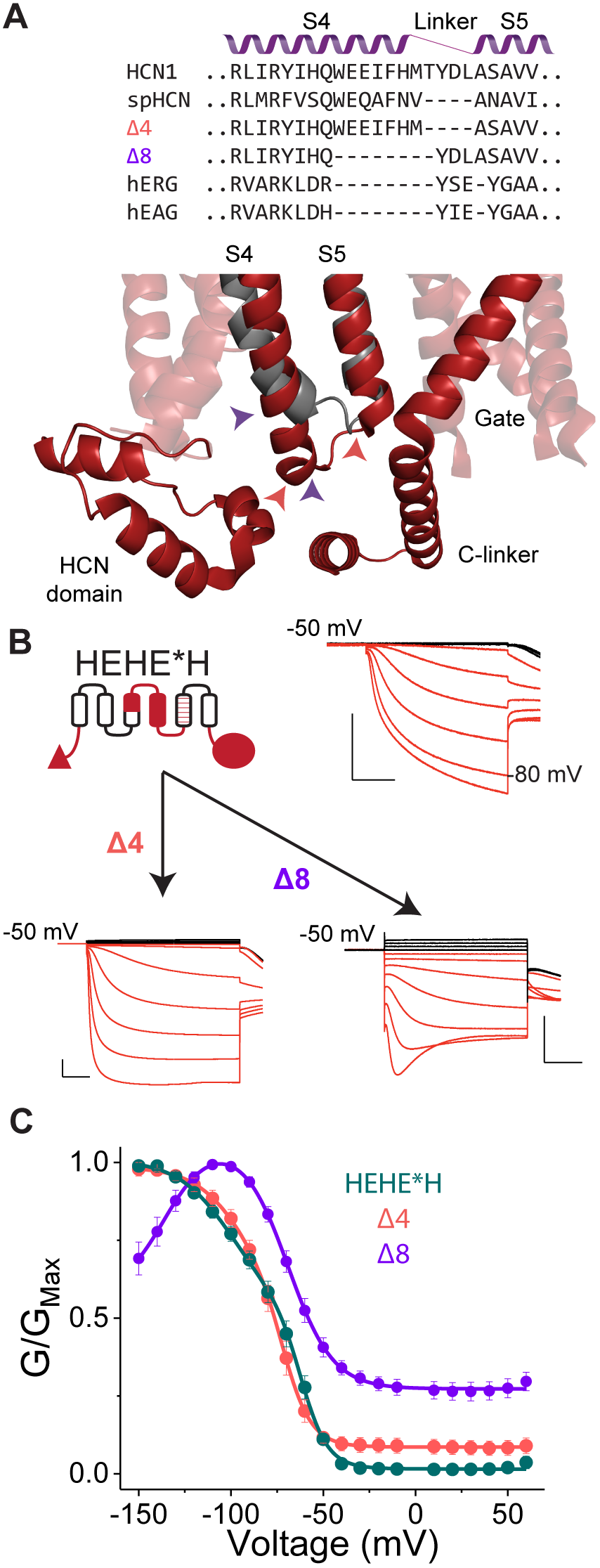
Long S4 Helix is not Required for Activation by Hyperpolarization. (A) Top: Sequence alignments for some HCN-and EAG-like channels in the S4-S5 helix-turn-helix region. The resulting sequences of the S4 deletion mutants generated in the background of the HEHE*H construct (Δ4 and Δ8) are also indicated. Bottom: HCN1 structure in the vicinity of S4-S5 turn of HCN1 (red) is overlaid with that of EAG1 (gray). Note the extended length of the HCN1 S4 and its close proximity to the HCN domain and C-linker, suggesting a potential for functionally critical interactions. Colored arrows indicate deletion boundaries in Δ4 (red) and Δ8 (purple) mutants. (B) Representative COVG current traces for the HEHE*H construct and its two deletion derivatives. Scale bars denote 2 μA (vertical) and 500 ms (horizontal). (C) GV curves for the parent HEHE*H and deletion mutants with error bars representing SEM for n= 4.

### Unified Voltage Gating Scheme for CNBD Family Channels

To evaluate whether the whole complement of phenotypes observed in this study can be quantitatively described by a simple unified gating model, we carried out kinetic simulations starting with an elementary scheme involving two distinct open states flanking a closed state at intermediate potentials (Figure 7 and Table S1). The underlying assumption throughout this exercise is that the gating scheme in this family of ion channels remains unchanged. The mutations (or swaps) change only the voltage-dependent rates of entry and exit from specific states. Figure 7A-B shows that the Janus phenotype observed in HHHEE and HHHEH chimeras can be easily described by this elementary model involving two distinct open states. The addition of the C-terminus stabilizes the hyperpolarization-dependent opening (O_H_) while simultaneously destabilizing the depolarization-dependent open state (O_D_). This is consistent with the notion that these two states are likely to be conformationally distinct.

**Figure 7.**
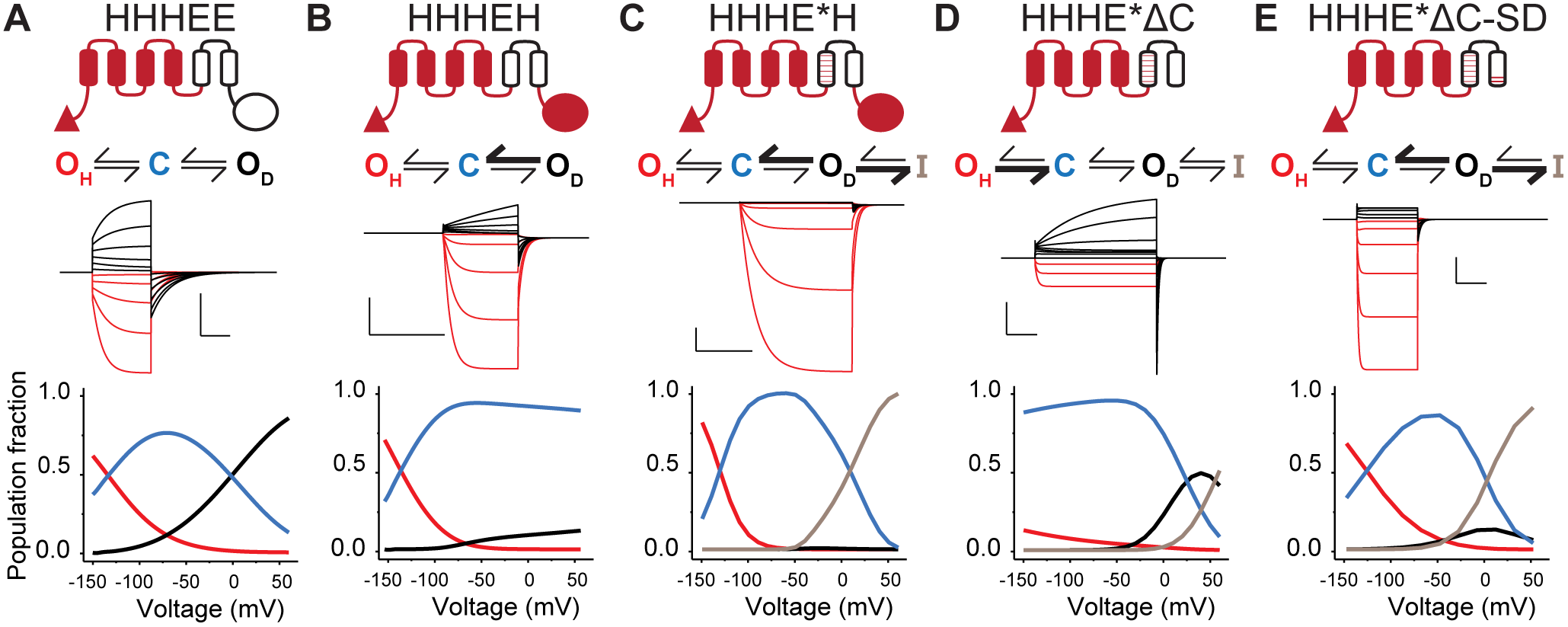
Unified Voltage Gating Scheme for CNBD Family of Channels. (A-E) Cartoon representations, models, simulated currents, and voltage dependencies of the population fractions of each state in the model. Bolded arrows indicate strengthened transitions. The population fraction curves are colored according to the models with the hyperpolarized open state in red, closed state in blue, depolarized open state in black, and inactivated state in gray. (F) Cartoon illustrating the proposed model of voltage-dependent gating applicable for our constructs and generally to all channels in the CNBD family. Cylinders represent S4 (blue), S5 helix (orange), and S6 helix (grey).

In order to account for the hooked tails in the mosaics with a reconstructed HCN1-like S4 and S5 interface, an inactivated state (I) was added to the depolarization-dependent open state (as described in the hERG gating model inWang et al., 1997). Parameterization of this expanded model by fitting them to current traces from HHHE*H shows that this expanded model best fits the data when the channels enter the inactivated state rapidly but their rate of entry to the depolarization dependent open state is relatively slow. Under these circumstances, the depolarization-dependent opening is obscured by rapid inactivation (Figure 7C). Comparing the population fraction plots for HHHEH and HHHE*H reveals that the main difference is that HHHE*H populates the inactive state at depolarized potentials, consistent with the notion that the primary effect of the interface is depleting the O_D_ state via inactivation (Figure 7B-C). Similar analysis of the HHHE*ΔC mosaic shows that its phenotype is fitted well by the same model simply by changing the rates to disfavor both the hyperpolarization-dependent opening as well as the entry into inactivated state (Figure 7D). This can be clearly seen by comparing the population fractions of HHHE*H and HHHE*ΔC: At negative potentials, there is an increased fraction of channels in the closed rather than O_H_ state for HHHE*ΔC, while at depolarized potentials there is an increase in O_D_ and decrease in the inactivated state. Finally, the behavior of HHHE*ΔC-SD can be recapitulated by adjusting the rates to increase stability of the hyperpolarization-dependent open state and inactivated state (Figure 7E). The population fraction plot of HHHE*ΔC-SD more closely resembles the HHHE*H mosaic rather that the parent HHHE*ΔC construct, suggesting the coupling of the hyperpolarization activation pathway is largely restored by these mutations (Figure 7C-E).

## DISCUSSION

In this study, we describe a hierarchical approach to determine how the various structural elements interact in a non-additive manner to control gating polarity in voltage-activated ion channels. Previous attempts of straightforward domain swaps (Cao et al., 1995; Poree et al., 2005; Riedelsberger et al., 2010) between outward-rectifying and inward-rectifying potassium channels failed to clarify this issue, largely due to a lack of high-resolution structures. One study, for instance, concluded that the elements involved in determining gating polarity are dispersed throughout the primary structure of the channel (Riedelsberger et al., 2010). The newly available structures of EAG1, hERG and HCN1 channels allow us to align them based on their secondary structural elements. These structural alignments along with sequence analysis of other orthologs in this family guided our choices of the junction points. Subsequently, by specifically focusing on recreating interaction pathways between the discrete structural entities, our refinement sheds new light on how coupling between these structural elements determine gating polarity.

According to the prevailing hypothesis, the hyperpolarization dependent opening in HCN channels is due to a reversal in the coupling machinery that results in pore gates being open when the voltage sensors are in down position, in contrast to the depolarization activated channels (Latorre et al., 2003; Mannikko et al., 2002). Our key finding described here is not compatible with this widely held view of “inverse coupling” which mandates that the same conformational change of the voltage sensor is used to gate the pore in one or the other direction. Our Janus chimeras and mosaics show that, in contrast to the previous studies (Cao et al., 1995; Li et al., 2008; Poree et al., 2005; Riedelsberger et al., 2010), both of these gating mechanisms can exist in the same channel. Although the ion permeation pathway is the same, our state-dependent blocker data argue that the conformation of the hyperpolarization activated open state is distinct from that of the depolarization-dependent opening.

The primary basis for HCN activation upon hyperpolarization is the specialized voltage response conferred by the extended S3b-S4 segment which inherently acts as a bipolar switch that drives channel opening in both directions. Our studies also show that the HCN voltage sensor movement alone is not sufficient to gate the EAG pore open upon hyperpolarization-other structural elements are necessary to stabilize the hyperpolarization gate in the open conformation. The C-terminus, which includes the C-linker and the CNBD, plays a crucial role by destabilizing the depolarization-dependent open state (O_D_) and favoring the hyperpolarization-dependent open state (O_H_) (Figure 7A-B, D). Additionally, specific residues in the vicinity of the pore gate strongly destabilize the depolarization-dependent open state. These residues, presumably, determine the local gate conformation. They must act independently of the C-terminal fragment because the HCN1Δ is still gated open by hyperpolarization (Wainger et al., 2001). Given that we do not have quantitative measures of coupling at this time, we cannot rule out that other as yet unidentified elements in the HCN1 pore further contribute to the stability of hyperpolarization-dependent open state.

Although the tight voltage sensor-pore interface observed in the structure of HCN channel is distinctive (Lee and MacKinnon, 2017), we find that the primary role of these interfacial interactions is to depopulate O_D_ by inducing the channels to rapidly enter into an inactivated state. Although this inactivation is reminiscent of hERG inactivation, the recently solved cryo-EM structure of hERG in the open state is missing this tight interface between S4 and S5 (Wang and MacKinnon, 2017). As the structure of the inactivated state is not known, further tests will be required to determine the role of this interface in hERG inactivation. Overall, our studies provide strong evidence that the hERG channels may serve as mechanistic link between the depolarization-activated and hyperpolarization-activated channels.

Given the strikingly long S4 helix and its close apposition to the S6 helix and cytoplasmic domains in the HCN1 structure, Lee and MacKinnon (2017)posited that the extra length of the HCN1 voltage sensor allows it to pin down the S6 gates in a closed position even when the voltage sensor is in the activated state. In contrast, the shorter S4 helices of the depolarization-activated channels such as EAG1 and hERG are unable to sterically prevent the opening of pore gates in the depolarized, upward conformation. Further, they propose that the extended length of S4 is also central to its role in the activation by hyperpolarization. Its downward movement upon further hyperpolarization presumably applies mechanical stress on the C-linker which is relieved by propagating a structural rearrangement to the gate-forming S6 helix.

Our findings indicate that such an explanation is incomplete. Indeed, a number of our chimeras contain full length S3b-S4 from HCN1 and yet are partially or exclusively activated by depolarization (Figure 1). Truncating the S4 to match the length of EAG S4 also does not eliminate activation upon hyperpolarization although it increases the basal leak through these channels. Thus, the long S4 by itself is not critical for closing the channel on depolarization or opening the channel on hyperpolarization.

If it is not the length of the S4 helix, then what enables the HCN voltage sensor to gate the pore upon hyperpolarization? Given the short length of the linker between the S4 and S5 segments, it is quite apparent that any significant downward movement of the S4 will be limited by S5. Therefore, one might anticipate that additional downward force upon hyperpolarization causes the S4 helix to undergo a conformational change that involves part of the S4 helix bending or unwinding. Such a conformational switch could disrupt S4-S5 packing and relieve the inhibition on the pore gates allowing them to open. Note that hyperpolarization-activated channels contain at least one serine residue, which is a known helix breaker (Ballesteros et al., 2000; Hall et al., 2009), in place of a hydrophobic residue in the middle of S4 helix (Figure S1). The notion that the HCN S4 helix undergoes a bending or twisting motion, rather than a large vertical displacement observed in canonical VGICs (Ahern and Horn, 2005; Chanda et al., 2005; Larsson et al., 1996), is also supported by cysteine accessibility studies on HCN (Bell et al., 2004; Vemana et al., 2004). These studies show that upon hyperpolarization, the S4 residues on the outside undergo a limited change in solvent accessibility whereas the accessibilities of those on the inside change drastically, consistent with widening of the inside facing crevice.

Although further testing will be necessary to evaluate this model of hyperpolarization gating, our “bipolar switch” model provides a unifying framework for understanding how the channels in the CNBD family give rise to such distinct phenotypes despite having a common architecture. Finally, these chimeras and mosaics greatly expand our existing repertoire of channel phenotypes and may serve as model systems for further structural and mechanistic analyses of electromechanical coupling.

## AUTHOR CONTRIBUTIONS

This work was designed by J.Cowgill, V.A. Klenchin, and B. Chanda. Constructs were primarily generated by V.A. Klenchin with assistance from J. Cowgill, C. Alvarez-Baron, D. Tewari, and A. Blair. Data was collected and analyzed by J. Cowgill, C. Alvarez-Baron, D. Tewari, and A. Blair. The manuscript was prepared by J. Cowgill, V.A. Klenchin, and B. Chanda with input from others.

## ACKNOWLEDGEMENTS

We would like to thank G.A. Robertson for providing hEAG1, C. Czajkowski for providing the pUNIV vector, B Santoro and S.A. Siegelbaum for providing mHCN1, M.C. Sanguinetti for providing mHCN2, and U.B. Kaupp for providing spHCN. We would also like to thank N. Nallappan for help generating chimeras and N. Nallappan, W. Stevens-Sostre, and T. Tsao for performing frog surgeries and providing oocytes. M.B. Jackson, L. Delemotte, and G.A. Robertson for their helpful comments and discussions. We also thank M. Kasimova and L. Delemotte for sharing unpublished results of their molecular dynamics simulations. This work was supported by funding from National Institutes of Health to B.C. (NS101723), J.B.C (T32 HL-07936-17) and C.A-B (T32 HL-07936-15), American Heart Association to C.A-B (17POST33411069), Romnes faculty fellowship to B.C. and Hilldale fellowships to A.B. The authors declare no competing interests.

## METHODS

### EXPERIMENTAL MODEL DETAILS

Oocytes from *Xenopus laevis* were isolated via surgery and digested using 0.8 mg/mL Collagenase II (Roche) for approximately one hour until the removal of the follicular layer. Oocytes were maintained in ND96 solution (96 mM NaCl, 2 mM KCl, 1 mM CaCl_2_, 1 mM MgCl_2_, 5 mM HEPES at pH 7.4) prior to injection then transferred to ND96 containing antibiotics (50 μg/mL of gentamicin and ciprofloxacin, 100 μg/mL tetracycline, penicillin, and streptomycin) and BSA (0.5 mg/mL) after injections.

### METHOD DETAILS

#### Alignments

Structural alignments were carried out by Pymol 1.8 (Schrodinger, 2015) using the default settings from the “super” command. Sequences for hHCN1 (Uniprot ID O60741), mHCN2 (O88703), spHCN (O76977), LKT1 (Q9LEG6), KAT1 (Q39128), SKOR (Q9M8S6), rEAG1 (Q63472), and hERG (Q12809) were first aligned by T-Coffee Expresso (Notredame et al., 2000) and then manually adjusted based on structural alignments.

#### Molecular Biology

Chimeras between mHCN1 (NCBI accession:NM_010408.3) and hEAG1 (NM_172362.2) were generated in pUNIV vector (Venkatachalan et al., 2007) as fusions with *X. laevis* codon-optimized mCherry gene at the 3’ end. Chimeragenesis was done by a QuikChange Cloning protocol that involves performing a standard QuikChange mutagenesis reaction in which a PCR-synthesized fragment of interest with terminal arms that anneal to the target plasmid is used in place of mutagenic oligonucleotides. The conditions and reagents used were exactly as previously described (Klenchin et al., 2011). Mosaic mutations were using similarly introduced by QuikChange Cloning using gBlock DNA fragment (Integrated DNA technologies). Point mutations were introduces by a QuikChange protocol using high fidelity PfuUltra II Fusion polymerase (Agilent) and a single mutagenic primer in the reaction. In all cases, the sequence of the entire ORF was verified by Sanger sequencing of both DNA strands. Chimeras with spHCN (NM_214564.1) or mHCN2 (NM_008226.2) were generated in the psGEM vector.

#### Oocyte microinjection

Oocytes were microinjected with 20-70 nL corresponding to 5-75 ng cRNA using a Nanoject II (Drummond Scientific). Injections were performed in ND96 solution without calcium chloride, then transferred to the post-injection solution described in the Experimental Model section.

#### Electrophysiology

Cut-open Vaseline gap (COVG) voltage clamp recordings were obtained at room temperature (22°C) with a CA-1B amplifier (Dagan) at a sampling rate of 10 kHz. Two-electrode voltage clamp (TEVC) recordings were obtained at room temperature with an OC-725C amplifier (Warner) at a sampling rate of 10 kHz. Thin-walled glass pipettes (World Precision Instruments) were used with tip resistances of 0.2-0.8 kΩ and filled with 3M KCl. Unless otherwise noted, external solutions contained 100 KOH, 5 NaOH, 20 HEPES, and 2 CaCl_2_. Internal solutions were comprised of 100 KOH, 5 NaOH, 10 HEPES, and 2 EGTA. For COVG experiments, oocytes were permeabilized using 0.3% saponin in internal solution and washed out prior to recording. All solutions were adjusted to pH 7.4 using methanesulfonic acid. All recordings were obtained with no leak subtraction, though capacitance compensation was used in COVG experiments to improve resolution of constructs with fast kinetics. Currents at the end of the test pulse were converted to conductance by dividing by the potential of the test pulse (assuming a reversal potential of 0 mV due to symmetrical recording solutions). Due to the inability to determine conductance at 0 mV in symmetrical solutions, this point is omitted from all GV curves. Conductance at the peak tail was determined by dividing the peak tail current amplitude by the potential at the tail pulse (generally -80 mV). Data in GV curves was fitted to a sum of two Boltzmann curves in Origin with the function f(x)= O_1_+((A_1_-O_2_)/(1+exp(k_1_(V-V_1_))) + O_2_+((A_2_-O_2_)/(1+exp(k_2_(V-V_2_))) where A_1_ and A_2_ represent the amplitudes, O_1_ and O_2_ represent the offsets, V_1_ and V_2_ represent the V_1/2_, and k_1_ and k_2_ represent the slope factors for two independent components. As many curves do not reach saturation, curves are primarily provided for visual reference.

#### MTSEA Block Experiments

MTSEA block was assessed on TEVC. Oocytes were perfused with external solution and given repeated test pulses every 30 seconds. Perfusion was halted prior to test pulse to avoid additional noise in the recording. When recordings stabilized, perfusion was switched to external solution containing 5 mM MTSEA and pulses continued. This solution was prepared fresh for each oocyte from a 1 M stock of MTSEA dissolved in external solution and kept on ice, which was prepared daily.

#### Astemizole Block Experiments

Astemizole block was assessed on COVG by repeating 3 second test pulses every 20 seconds. External solution was exchanged in between pulses. Some oocytes showed current rundown following initial solution exchange. Once recordings were stable over at least 3 consecutive pulses, 50 μM astemizole was applied. After two pulses in the presence of the astemizole solution, external solution was exchanged back to the original solution (see Electrophysiology section). Pulses were continued until recordings stabilized.

#### Model Simulations

Kinetic models were built and simulated using the software KineticModelBuilder Version 2.0 described previously (Goldschen-Ohm et al., 2014). All rates were allowed to vary between 0 and 1000 s^-1^ while charge was limited to 0 to 3 e^-^. Initial values for the rate and charge of each step were set to 1 and then current traces were fit using the EigenSolver mode until convergence. Weighting was used to better recapitulate the kinetics of the process and avoid bias of over-fitting steady state. This was accomplished by weighting the fit immediately following the test and tail pulse 10 fold compared to steady state. This weight decayed exponentially back to baseline with a time constant of 10 ms. Population fractions at “steady state” were determined by the occupancy of each state in the model over the last 10 ms of the test pulse. Briefly, the state occupancy was determined by numerical solution of the transition matrices within the model as described previously (Colquhoun and Hawkes, 1995; Goldschen-Ohm et al., 2014).

### QUANTIFICATION AND STATISTICAL ANALYSIS

Clampfit (Molecular Devices) was used to quantitate currents at steady state. Origin was used to fit data points to a sum of two Boltzmann curves. Kinetic Model Builder was used for all kinetic simulations. Pymol was used for all structural analyses. Throughout the paper, n is used to denote the number of oocytes tested in each experiment as indicated in each figure legend.

## SUPPLEMENTAL FIGURE LEGENDS

**Figure S1.**
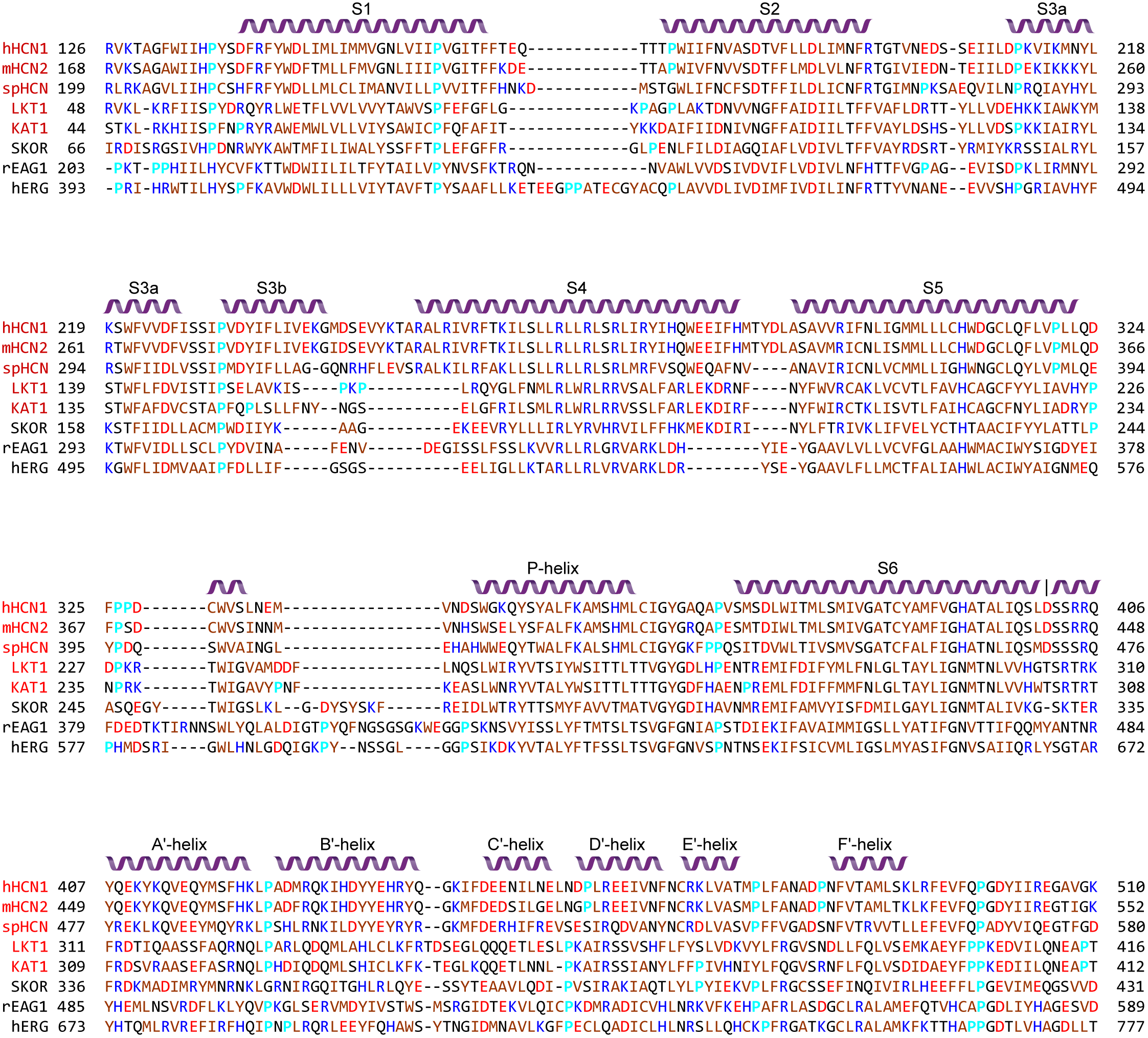
Sequence Alignment of Voltage-gated Ion Channels from CNBD Family. Sequence Alignment of the transmembrane segments of some well-studied voltage-gated channels in the CNBD family from mammals (human HCN1, mouse HCN2, rat EAG1, human hERG), invertebrates (sea urchin spHCN) and plants (tomato LKT1, arabidopsis KAT1 and SKOR). The channels activated by hyperpolarization are shown in red and those activated by depolarization are in black. The sequences were first aligned by T-Coffee Expresso (Notredame et al., 2000) and then manually adjusted to match experimentally-determined structures of hHCN1, rEAG1 and hERG (PDB entries 5U6O, 5K7L and 5VA1). Helical structures of hHCN1 are indicated by purple ribbons. The coloring scheme used divides residues into five groups: charged acidic, charged basic, hydrophobic/non-polar, uncharged polar and prolines.

**Figure S2.**
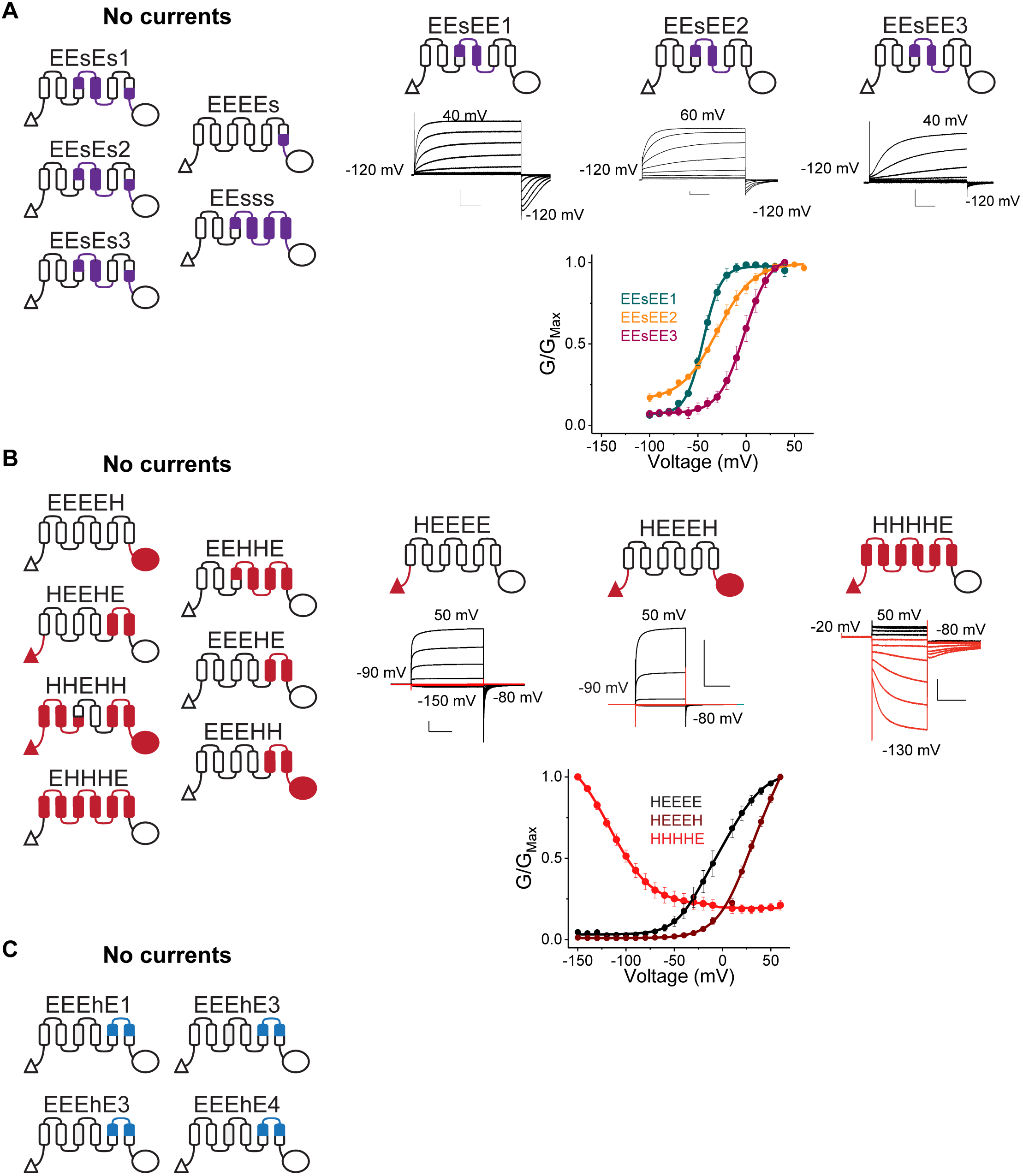
Initial Chimeras Tested. (A-C) Preliminary scanning for functional chimeras between hEAG1 and spHCN (A), mHCN1 (B), and mHCN2 (C). Cartoon representation of the constructs that did not display detectable currents are shown on the left. Functional constructs are shown on the right together with representative traces from two-electrode voltage clamp (for constructs in A) or cut-open voltage clamp (for constructs in B) and GV curves with errors representing SEM for n=3 (for constructs in A) or 4 (for constructs in B).

**Figure S3.**
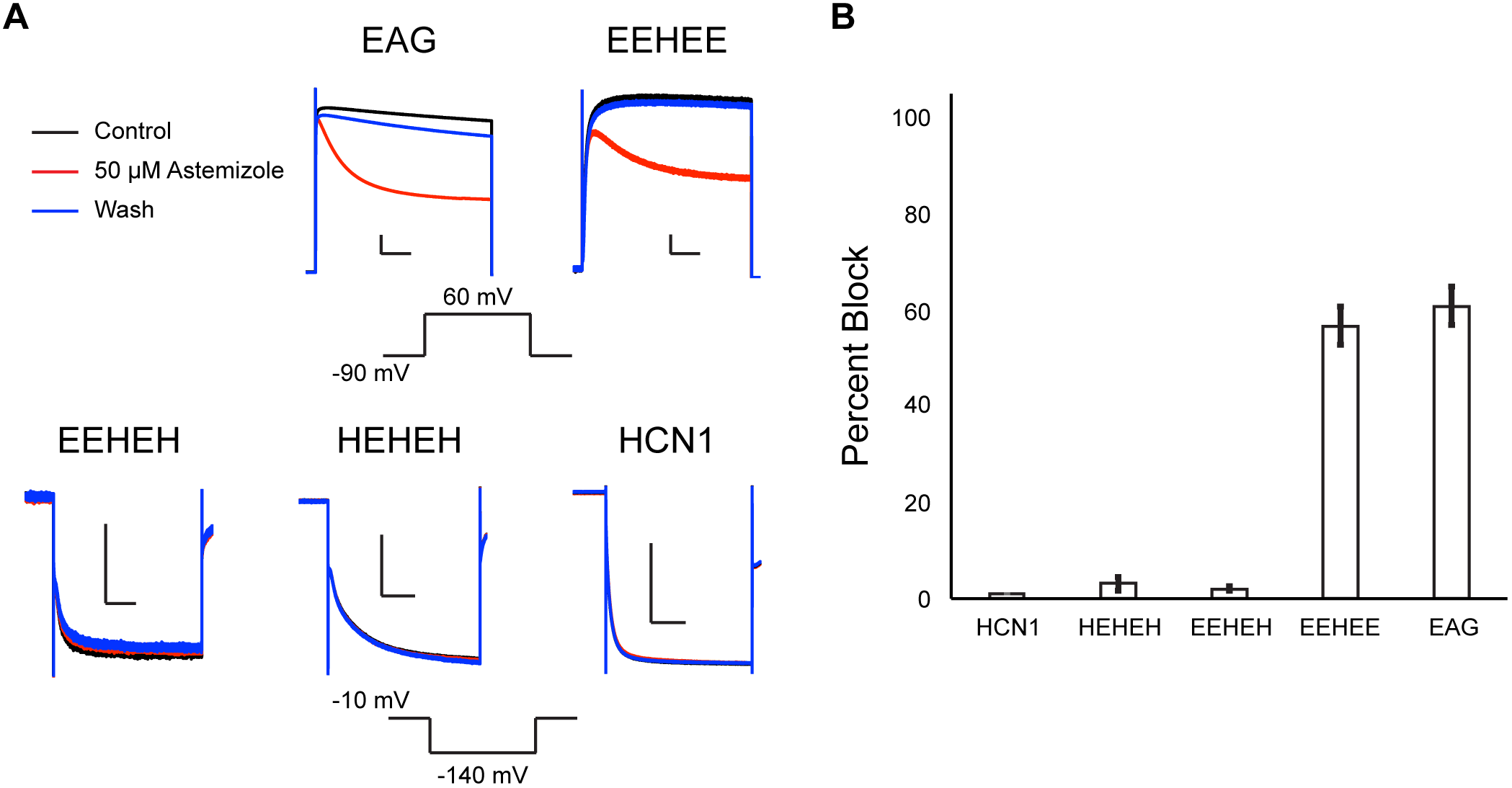
Hyperpolarization-activated Currents Are Not Blocked by Astemizole. (A) Representative current traces before (black), after application of 50 μM astemizole (red), and after wash (blue). (B) Percent of current blocked for each construct shown in panel A calculated based on the decrease in observed current at the end of the test pulse before and after application of astemizole. Error bars represent SEM for n=3.

**Figure S4.**
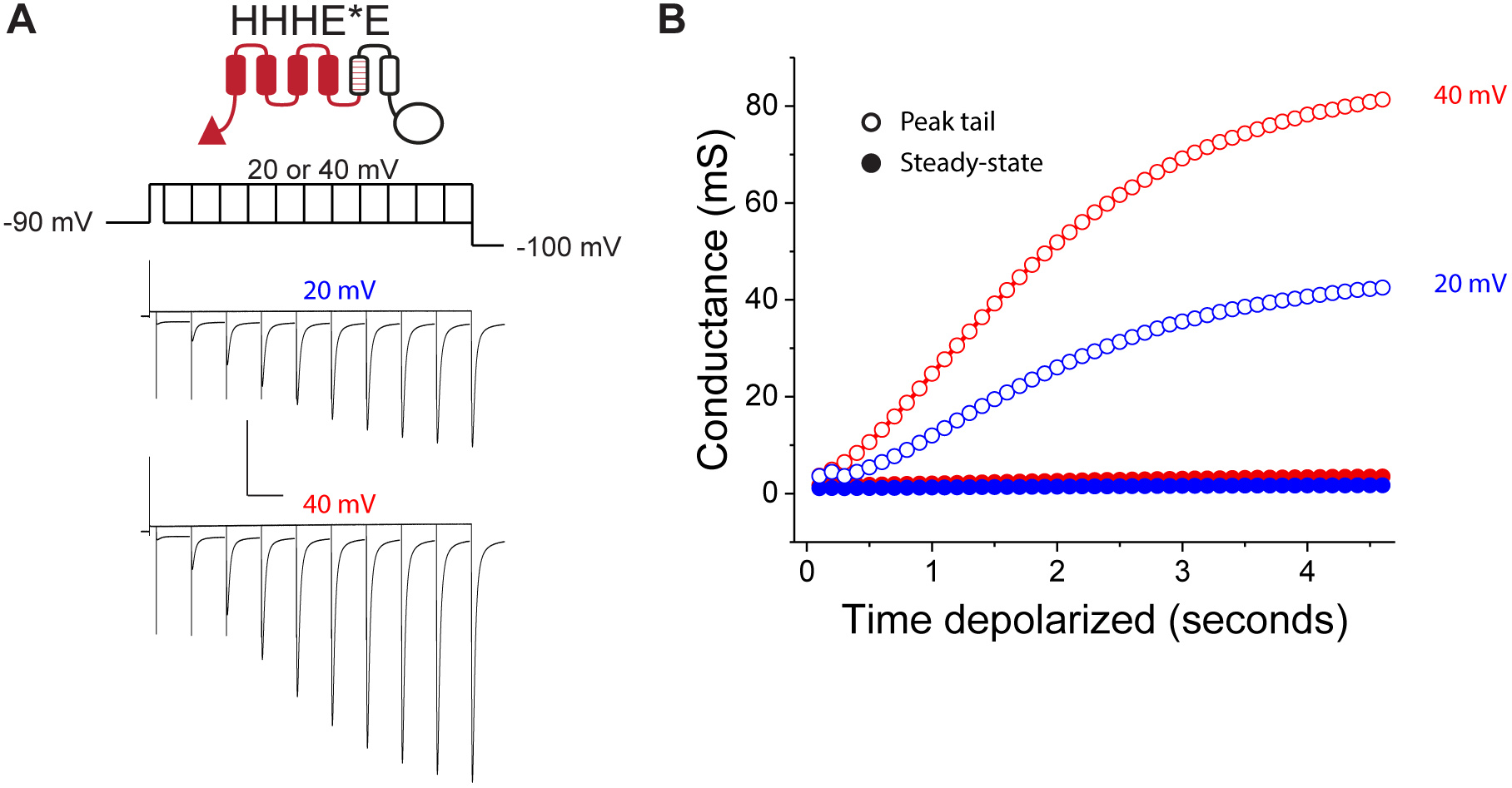
Activation Kinetics in the HHHE*H Construct. (A) Envelope of tails protocol under symmetric solutions (100 mM K^+^_internal_/100 mM K^+^_external_) in cut open voltage clamp showing the growth of tail currents elicited by depolarizing pulses of increasing length to 20 mV or 40 mV. (B) Conductance plotted as a function of time at steady-state (filled circles) or peak tail (open circles) from depolarizing pulses to 20 mV (blue) or 40 mV (red).

**Figure S5.**
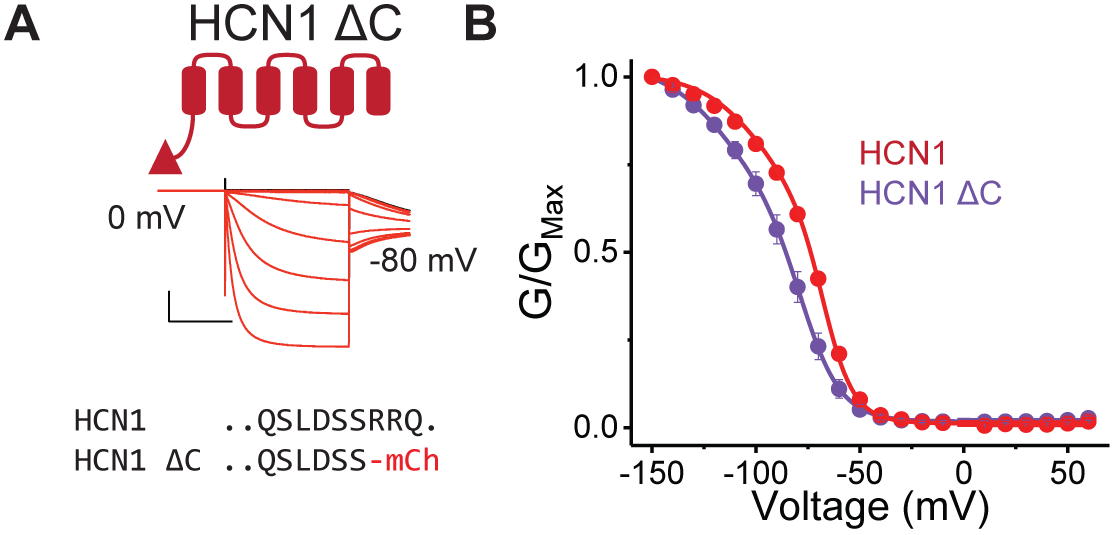
HCN1 is Minimally Affected by the Removal of its C-terminus. (A) Cartoon representation, sequence, and representative current traces for the C-terminal deletion of mHCN1 fused to mCherry. (B) GV curves for the wild type mHCN1 fused to mCherry and its C-terminal deletion construct depicted in (A). Experimental conditions were identical to those used to generate data show in Figure 1. Error bars represent SEM for n=5.

**Figure S6.**
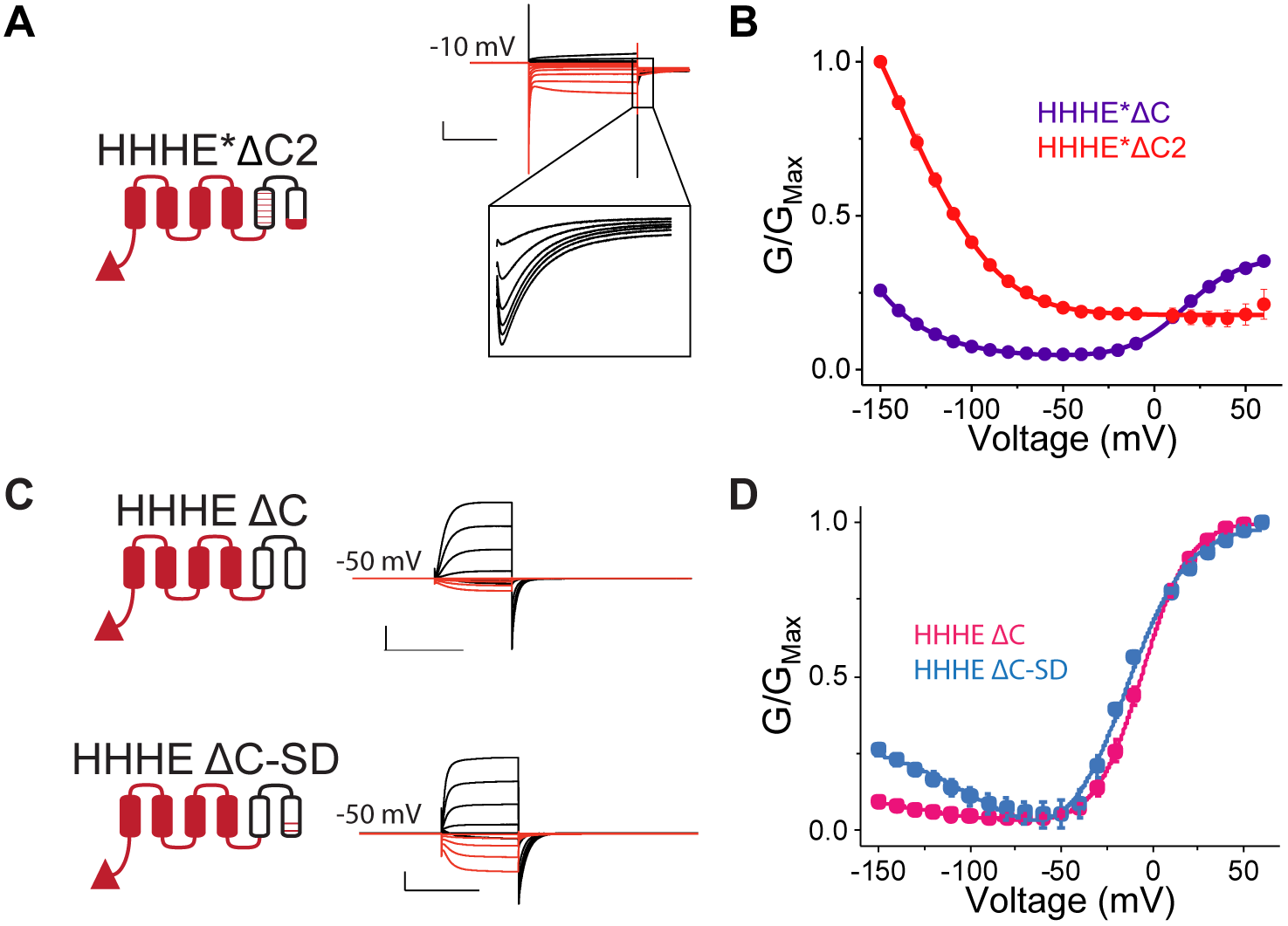
Residues Near the Channel Gate Influence Gating Polarity in the Absence of C-terminus. (A) Cartoon representation, sequence, and representative current traces for the C-terminal deletion mosaic that contains last eight HCN1-specific residues immediately following the gating glutamine residue. (B) Comparison of the GV curves between the original truncation construct and its HHHE*ΔC2 variant depicted in (A). (C) Cartoon representation and representative current traces for the HHHEΔC constructs tghat have same truncation as in HHHE*ΔC (see Figure 5D) but lack mosaic mutations on S5. Shown are data for the “wild type” HHHEΔC and its double mutant derivative with Q540S/Y542D mutations that are the same as found in the equivalent mosaic construct HHHE*ΔC (see Figure 5D). (D) GV curves for HHHEΔC and HHHEΔC-SD with error bars representing SEM for n= 3 (HHHEΔC-SD) or 4 (HHHEΔC).

**Table S1.**
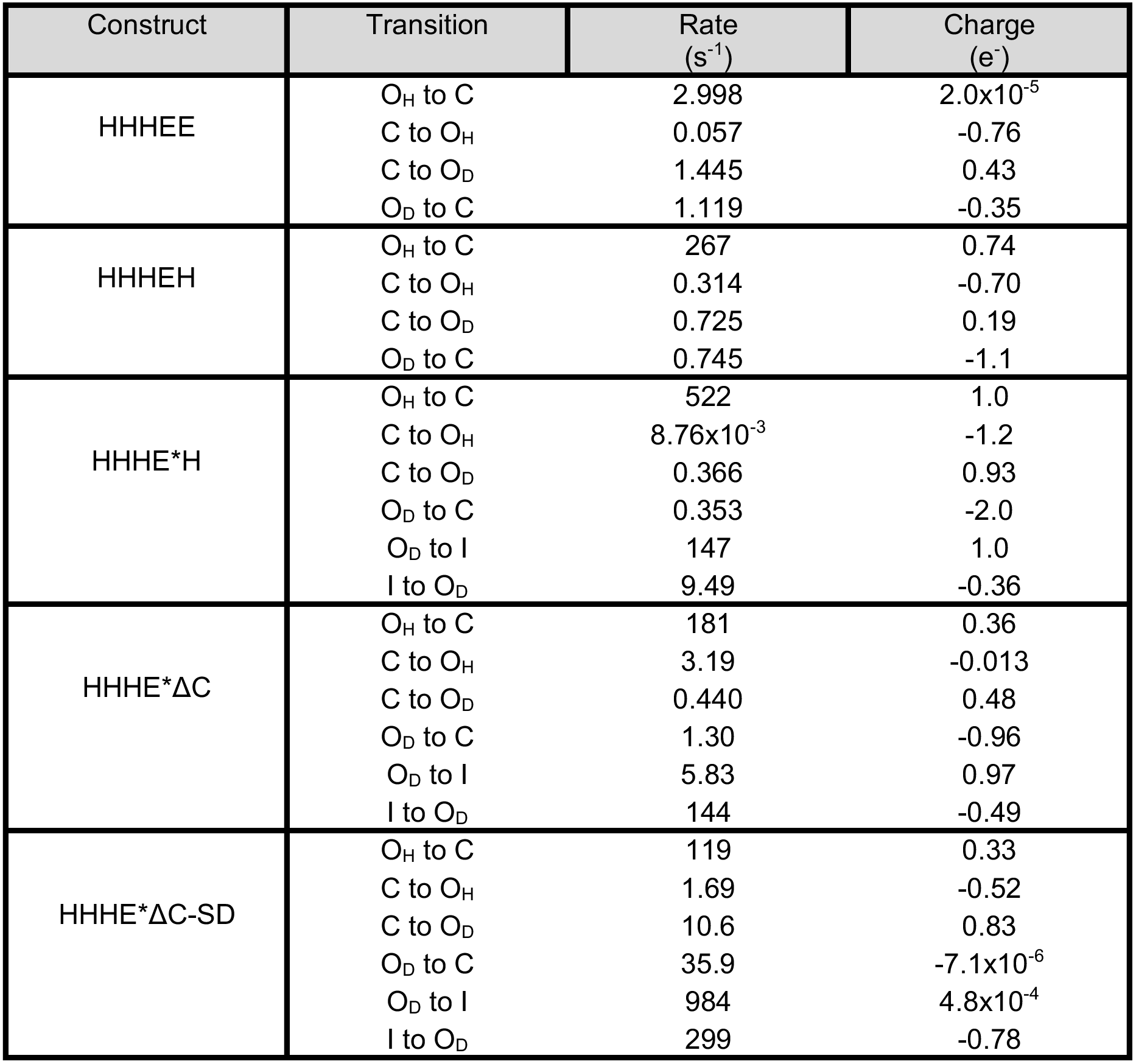
Related to Figure 6 Parameters for kinetic models used in simulations

